# Precise removal of *Calm1* long 3′ UTR isoform by CRISPR-Cas9 genome editing impairs dorsal root ganglion development in mice

**DOI:** 10.1101/553990

**Authors:** Hannah N. Gruner, Bongmin Bae, Maebh Lynch, Daniel Oliver, Kevin So, Grant S. Mastick, Wei Yan, Pedro Miura

**Author notes:** Corresponding author: Pedro Miura Department of Biology, University of Nevada, Reno. 1664 N. Virginia St, Reno, NV 89557, USA. these authors should be considered co-first authors (Hannah N. Gruner, Bongmin Bae). The authors declare no competing financial interests.

## Abstract

Most mammalian genes are subject to Alternative cleavage and PolyAdenylation (APA), which most commonly leads to the expression of two alternative length 3′ UTR isoforms. Long 3′ UTR variants are preferentially expressed in neuron-enriched tissues of metazoans. Their functional relevance *in vivo* is largely unknown. *Calmodulin 1* (*Calm1)* is a key integrator of calcium signaling that is required for correct neural development that generates short (*Calm1-S*) and long (*Calm1-L*) 3′ UTR mRNA isoforms via APA. We found *Calm1-L* expression to be largely restricted to neuronal tissues in mice with pronounced expression in the Dorsal Root Ganglion (DRG), whereas *Calm1-S* was more broadly expressed. Within DRG neurons, *Calm1-S* expression was detected in DRG axons, whereas *Calm1-L* was mostly restricted to soma. To study the *in vivo* function of *Calm1-L* we devised a CRISPR-Cas9 genome editing approach to generate mice that lack *Calm1-L* expression but maintain normal expression of *Calm1-S*. Three mutant mice lines were established that harbored different deletions, all of which successfully eliminated *Calm1-L*. Embryos lacking *Calm1-L* were found to have disorganized DRG axon and cell body migration. Explant DRG cultures from these mice were stimulated with Nerve Growth Factor to promote axon outgrowth and found to exhibit a dramatic increase in axon fasciculation. While overall CaM protein levels were found to be unaltered by long 3′ UTR loss, RNA stability assays in primary neurons revealed that *Calm1-L* is less stable than *Calm1-S*. Together, these results demonstrate the requirement for *Calm1-L* in DRG development and establish a widely applicable genome-editing strategy for generating long 3′ UTR isoform mutant mice.

## Introduction

Alternative cleavage and PolyAdenylation (APA) is the process by which a pre-mRNA is cleaved at two or more polyadenylation sites (polyA sites). This commonly results in two mRNAs with the same protein coding sequence but different length 3′ UTRs. APA is pervasive, occurring in ~51-79% of mammalian genes (Hoque et al. 2013; Lianoglou et al. 2013). As 3′ UTRs are major targets for post-transcriptional regulation via microRNAs (miRNAs) and RNA Binding Proteins (RBPs), harboring a longer 3′ UTRs can confer additional regulatory opportunities for transcripts (Sandberg et al. 2008; Mayr and Bartel 2009). Long 3′ UTRs impact translation in a cell context-specific manner (Floor and Doudna 2016; Blair et al. 2017), and elements located in alternative 3′ UTRs can influence mRNA localization in neurons (Taliaferro et al. 2016; Tushev et al. 2018b). Thousands of genes in mouse and human were recently found to express novel long 3′ UTR isoforms in brain tissues (Miura et al. 2013). Some of these genes were validated to express short 3′ UTRs isoforms across many tissues, whereas the alternative long 3′ UTR isoforms were found to be abundant only in brain tissues. The impact of neural-enriched, long 3′ UTR isoforms on nervous system function is poorly understood, despite the large number of genes shown to be affected in *Drosophila*, Zebrafish, mice, and humans (Smibert et al. 2012; Ulitsky et al. 2012; Miura et al. 2013).

Multiple studies have examined the functions of 3′ UTRs *in vivo* through generation of targeted deletions in mice. These studies have underscored the relevance of the 3’ UTR for controlling mRNA localization to axons and dendrites (Miller et al. 2002; Andreassi et al. 2010; Perry et al. 2012; Terenzio et al. 2018). Only a handful of studies have investigated *in vivo* functions of alternative long 3′ UTR transcripts. Recently, a lentiviral shRNA approach was used to knock down the long 3′ UTR isoform of *Rac1* in mouse primary cortical neurons, revealing defects in dendrite outgrowth (Braz et al. 2017). A genetic approach was implemented in mice to abolish the BDNF long 3′ UTR isoform while maintaining expression of its short mRNA counterpart via the insertion of tandem SV40 polyA sites downstream of the proximal polyadenylation signal. These mice displayed synaptic defects and hyperphagic obesity, ostensibly due to compromised mRNA localization to dendrites and impaired translational control (An et al. 2008; Liao et al. 2012). The recent advent of CRISPR-Cas9 genome editing has revolutionized the speed and efficiency of generating deletion mouse strains (Wang et al. 2013). This presents an exciting new opportunity for rapidly generating 3′ UTR isoform-specific knockout mice. To date, successful generation of a long 3′ UTR knockout mouse using CRISPR-Cas9 has not been reported.

Calmodulin (CaM) is the primary calcium sensor in the cell (Means and Dedman 1980; Yamniuk and Vogel 2004; Sorensen et al. 2013). CaM is expressed ubiquitously but is particularly abundant in the nervous system (Kakiuchi et al. 1982; Ikeshima et al. 1993). There are three *Calmodulin* genes in mammals– *Calm1*, *Calm2*, and *Calm3*. These share an identical amino acid coding sequence, but possess unique 5’ and 3′ UTRs (SenGupta et al. 1987; Fischer et al. 1988), suggesting differences in their regulation might be conferred at the post-transcriptional level. The existence of alternative short and long 3′ UTR mRNA isoforms of the *Calmodulin 1* (*Calm1*) gene have been known for several decades (SenGupta et al. 1987; Nojima 1989; Ni et al. 1992; Ikeshima et al. 1993). These isoforms include mRNAs with a short 0.9 kb 3′ UTR (*Calm1-S*) and one with a long 3.4 kb 3′ UTR (*Calm1-L*). The functional significance of these alternative 3′ UTR isoforms is unknown.

Previous work has shown that altering CaM levels disrupts neuronal development in *Drosophila* and rodents (Vanberkum and Goodman 1995; Fritz and VanBerkum 2000; Kim et al. 2001; Kobayashi et al. 2015; Wang et al. 2015). CaM plays a role in guiding axon projections to create connections with other neurons or tissues (Vanberkum and Goodman 1995; Fritz and VanBerkum 2000; Kim et al. 2001; Kobayashi et al. 2015). Functions in the nervous system specifically for *Calm1* have been described. Notably, targeted knockdown of *Calm1*, but not *Calm2* or *Calm3,* was found to cause major migration defects in developing hindbrain neurons (Kobayashi et al. 2015). *Calm1* mRNA has been detected in cultured rat embryonic DRG axons, where they undergo local translation to promote axon outgrowth (Wang et al. 2015). In these studies, whether *Calm1-L* specifically plays a role in neuronal development was not addressed.

Here, we utilized CRISPR-Cas9 to generate the first mouse lines lacking a long 3′ UTR mRNA isoform from the genome while maintaining normal expression of its short 3‱ UTR counterpart. Phenotypic analysis of mutant embryos revealed disorganized DRG axon and cell body positioning. Mutant DRG explants grown in culture exhibited increased fasciculation of axons upon NGF stimulation. This novel deletion methodology has thus uncovered a neurodevelopmental phenotype caused by removing a long 3′ UTR isoform in mice.

## Results

### Calm1-L expression is enriched in neural tissues

To determine the relative expression of short and long *Calm1* isoforms among mouse tissues, we performed Northern analysis– an approach uniquely suited to report on the expression of alternative 3′ UTR isoforms. Northern blots performed using a universal probe (“uni”) that detects both short and long *Calm1* isoforms revealed the presence of both *Calm1-L* and *Calm1-S* among an array of adult tissues. The cortex showed greater enrichment of *Calm1-L* versus *Calm1-S* compared to non-brain tissues, as had been previously observed (Ikeshima et al. 1993; Miura et al. 2013) (Fig. 1a,b). For quantification, we determined the ratio of long isoform to short isoform band intensity (L/S), and set the liver L/S ratio to 1.0 (Fig. 1b). The L/S ratio was 11.8-fold greater in cortex relative to liver (Fig. 1b). Recent studies have shown that some 3′ UTRs can be cleaved to generate stable 3′ UTR transcripts that are separated from the protein coding portion of the mRNA (Kocabas et al. 2015; Malka et al. 2017; Sudmant et al. 2018). The blot was stripped and reprobed with a probe specific for *Calm1-L* (“ext”). This revealed only one band consistent with the size of *Calm1-L* (Fig. 1a,c). Lower molecular weight bands were not observed, suggesting that if cleaved 3′ UTR *Calm1* transcripts are generated from the *Calm1* long 3′ UTR, they are expressed at levels lower than the detection limits of this Northern analysis.

**Figure 1.**
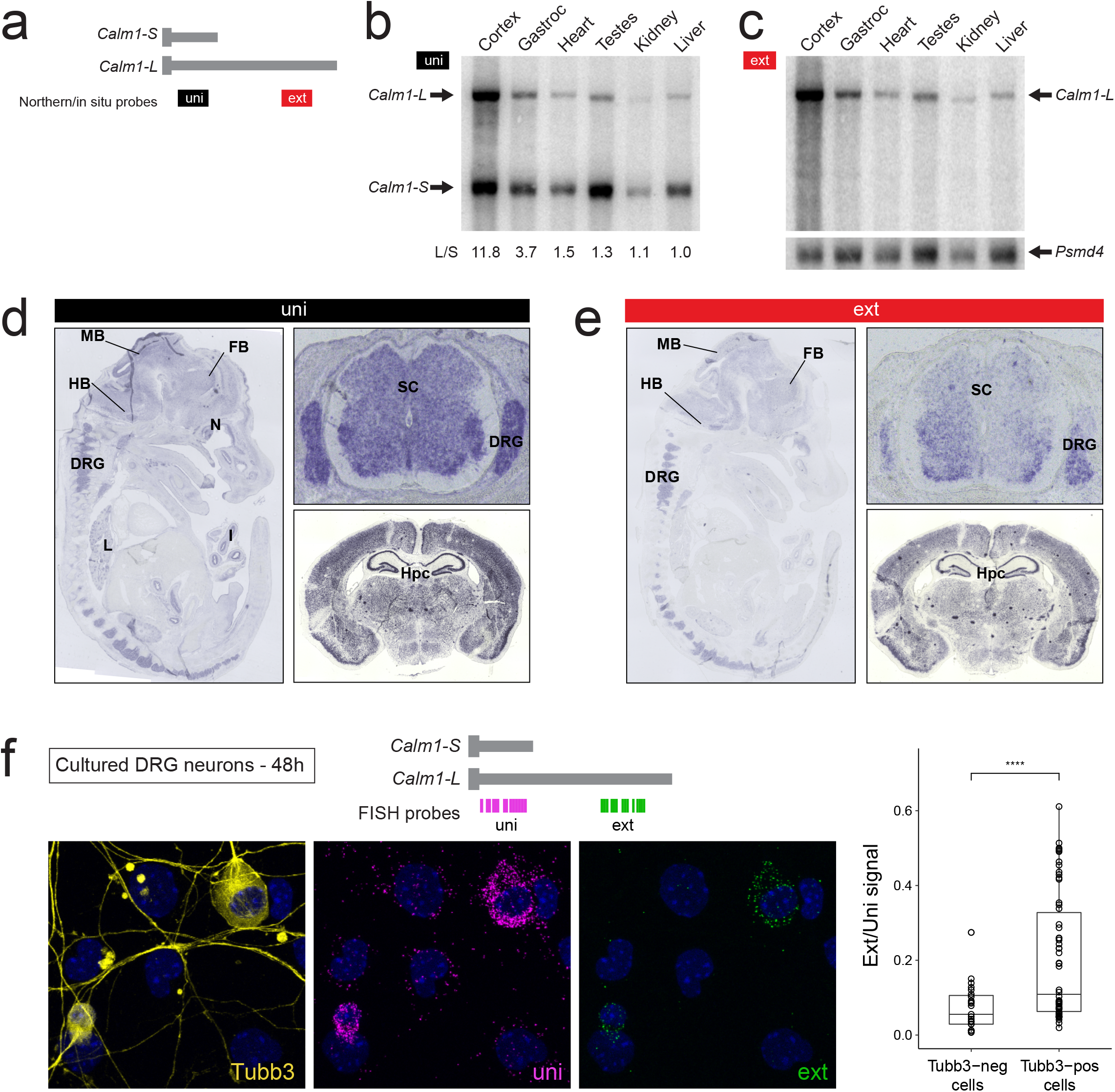
*Calm1* long 3′ UTR isoform (*Calm1-L*) is enriched in neural tissues. **(a)** Diagram showing location of Northern blot and DIG *in situ* probes to detect either both isoforms (uni) or specifically the long isoform (ext) of *Calm1*. **(b)** Northern blot of adult mouse tissues probed with “uni” probe reveals the presence of *Calm1-S* (bottom arrow) and *Calm1-L* (top arrow). The ratio of *Calm1-L* normalized to *Calm1-S* (L/S) is shown with liver ratio set to 1.0. **(c)** Northern blot performed with “ext” probe on the same blot as in (b) is shown. Note that a third, shorter 3′ UTR isoform was previously characterized in mouse testis (Ikeshima et al. 1993) but our probe design precludes detection of this transcript. Bottom blot shows *Psmd4* bands on the same blot as loading control. **(d)** Left: DIG *in situ* hybridization of E13.5 mice using uni probe shows signal in the forebrain (FB), midbrain (MB), hindbrain (HB), dorsal root ganglion (DRG), nasal epithelium (N), lung (L), and intestinal epithelium (I). Top right: Transverse section showing expression in spinal cord (SC) and DRG. Bottom right: P14 brain coronal section showing enrichment in hippocampus (Hpc). **(e)** DIG *in situ* performed with the ext probe shows a neuronal-specific expression pattern of *Calm1-L*. **(f)** Fluorescence *in situ* hybridization (FISH) assay coupled with anti-β3 Tubulin staining in cultured DRGs suggests *Calm1-L* is preferentially expressed in DRG neurons. Probe locations shown on top. Right panel shows boxplot of ext/uni signal (n = 23 Tubb3-neg and n = 58 Tubb3-pos cells, 4 independent culture preparations. t-test, ****: p < = 0.0001).

We next performed *in situ* hybridization using Digoxigenin-labeled probes in embryonic day 13.5 (E13.5) mice to determine the spatial expression pattern of *Calm1* isoforms in embryos. Using a *Calm1* universal probe, we observed strong signal in the brain, spinal cord, and dorsal root ganglion (DRG) (Fig. 1d). Signal was also observed in the nasal epithelium, lung, and intestinal epithelium. In contrast, *in situ* hybridization performed with the *Calm1* extension probe showed staining largely restricted to the brain, spinal cord, and DRG (Fig. 1e). The particularly strong extension probe signal in the DRG suggested to us that *Calm1-L* might have a biological role in the developing DRG.

The expression level of *Calm1-S* and *Calm1-L* was analyzed in cultured DRGs using RNAscope^®^ Fluorescence In Situ Hybridization (FISH). Co-staining with anti-*β* 3 Tubulin (Tubb3) was used to identify neurons. Quantification of universal and extension signals indicated that both isoforms of *Calm1* are abundantly expressed in Tubb3-positive cells. The ratio of extension to universal signal was found to be significantly higher in Tubb3-positive cells relative to Tubb3-negative cells (Fig.1f). This suggests that *Calm1-L* is preferentially expressed in neurons versus other cell types in the DRG.

### Calm1-S is more axonally iocalized than Calm1-L

Given earlier work showing that elements in the long 3′ UTRs of *importin β1* (Perry et al. 2012) and *Impa1* (Andreassi et al. 2010) are involved in mRNA localization to PNS axons, we hypothesized that *Calm1-L* might be specifically localized to axons. To test this hypothesis, we cultured dissociated E13.5 DRG neurons in compartmentalized chambers for two days and performed a FISH assay using universal and extension probes (Fig. 2a-c). Localization of *Calm1* transcripts was inspected along with coimmunostaining for β3 Tubulin in proximal and distal axons (Fig. 2b,c). In agreement with previous studies that showed *Calm1* expression in sensory neuron axons using FISH, microarray, and RNA-Seq analysis of axonal transcriptomes (Zivraj et al. 2010; Gumy et al. 2011; Wang et al. 2015; Preitner et al. 2016), we found that the universal probe detecting all *Calm1* isoforms revealed signal both in soma and proximal axons (Fig. 2b’; arrow heads in axons). Little to no signal was detected in distal axon compartment that overlapped with β3 Tubulin staining (Fig. 2c’) in 2 days culture. In contrast, there was extension signal indicating expression of *Calm1-L* in soma (Fig. 2b’’), but little to no signal in axons (Fig. 2b’’,c’’). Quantification of axonal punctae detected by the two probes in 2 days culture revealed a significant enrichment of universal signal over extension signal in DRG axons (Fig. 2d). This result demonstrates that *Calm1-L* is unlikely to harbor additional mRNA localization elements compared to *Calm1-S*.

**Figure 2.**
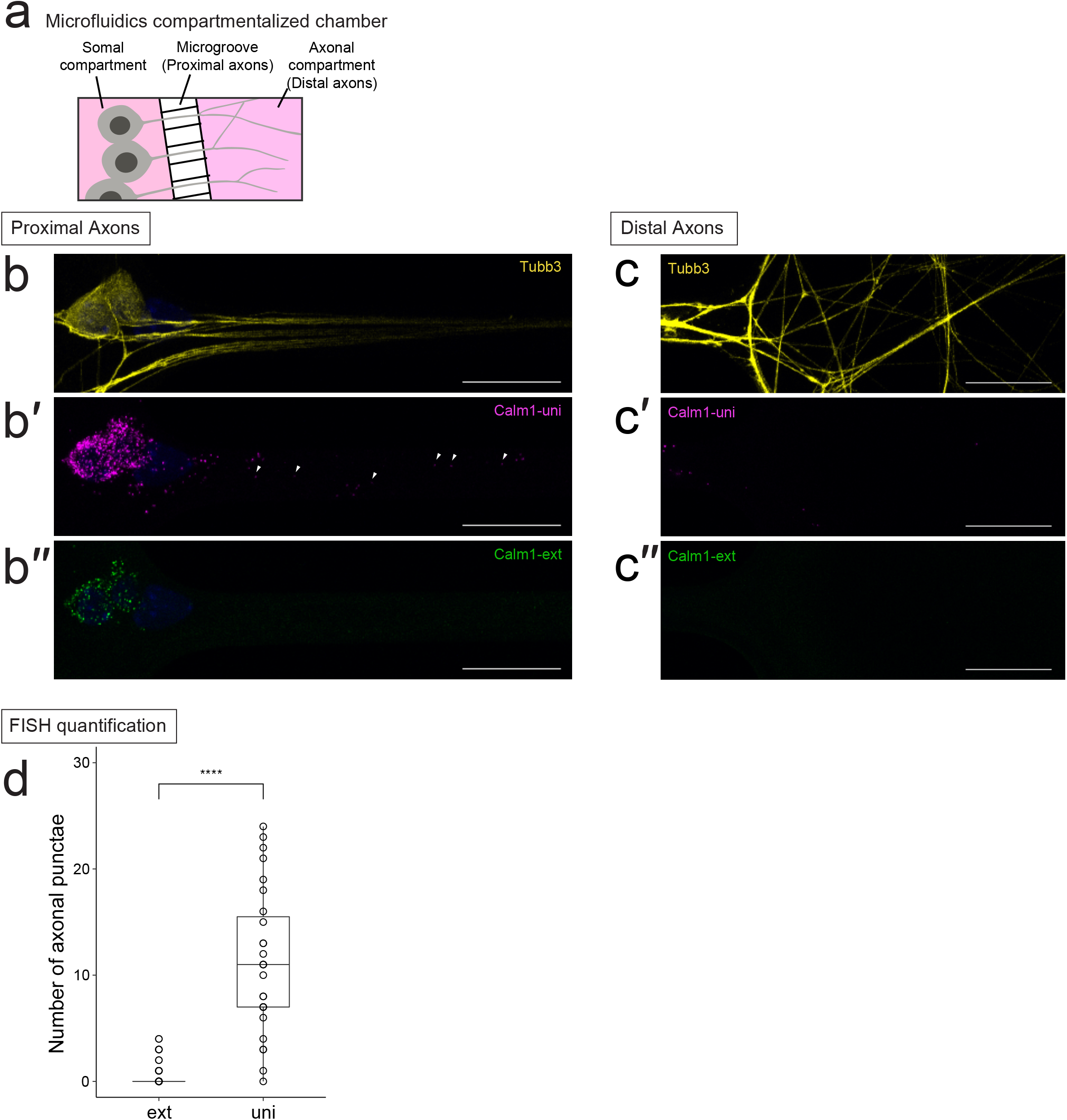
Localization of *Calm1 3′* UTR isoforms in DRG neurons. **(a)** Dissociated E13.5 DRG neurons were cultured in compartmentalized chambers for 48 hours to separate soma and axons. (**b-b″**) FISH coupled with β3 Tubulin (Tubb3) immunostaining to visualize *Calm1* isoforms in proximal DRG axons. Uni probe detects all *Calm1* isoforms whereas ext probe exclusively detects *Calm1-L*. Uni signals were detected in soma and axons. Ext signals were detected in soma whereas little to no signal was found in axons. (**c-c″**) In the distal axon compartment *Calm1* uni or ext punctae were not detected that overlapped with β3 Tubulin staining. Scale bar = 25 μm **(d)** Number of axonal punctae was counted for ext and uni signals (n = 30 cells, 2 independent culture preparations. t-test, ****: p <= 0.0001).

### Generation of Calm1 long 3′ UTR deletion mice using CRISPR-Cas9

To investigate the functional role of *Calm1-L in vivo* we employed CRISPR-Cas9 genome editing to generate a series of mouse lines that lacked *Calm1-L*. We wanted to generate a mutant line which did not harbor any foreign DNA sequences to promote usage of the proximal polyadenylation site over the distal polyadenylation site. We also wanted to completely abolish *Calm1-L* expression while maintaining normal expression levels of *Calm1-S*. We designed 6 guide RNAs (gRNAs) targeting the *Calm1* locus that we anticipated would result in three different deletions (Fig. 3a).

**Figure 3.**
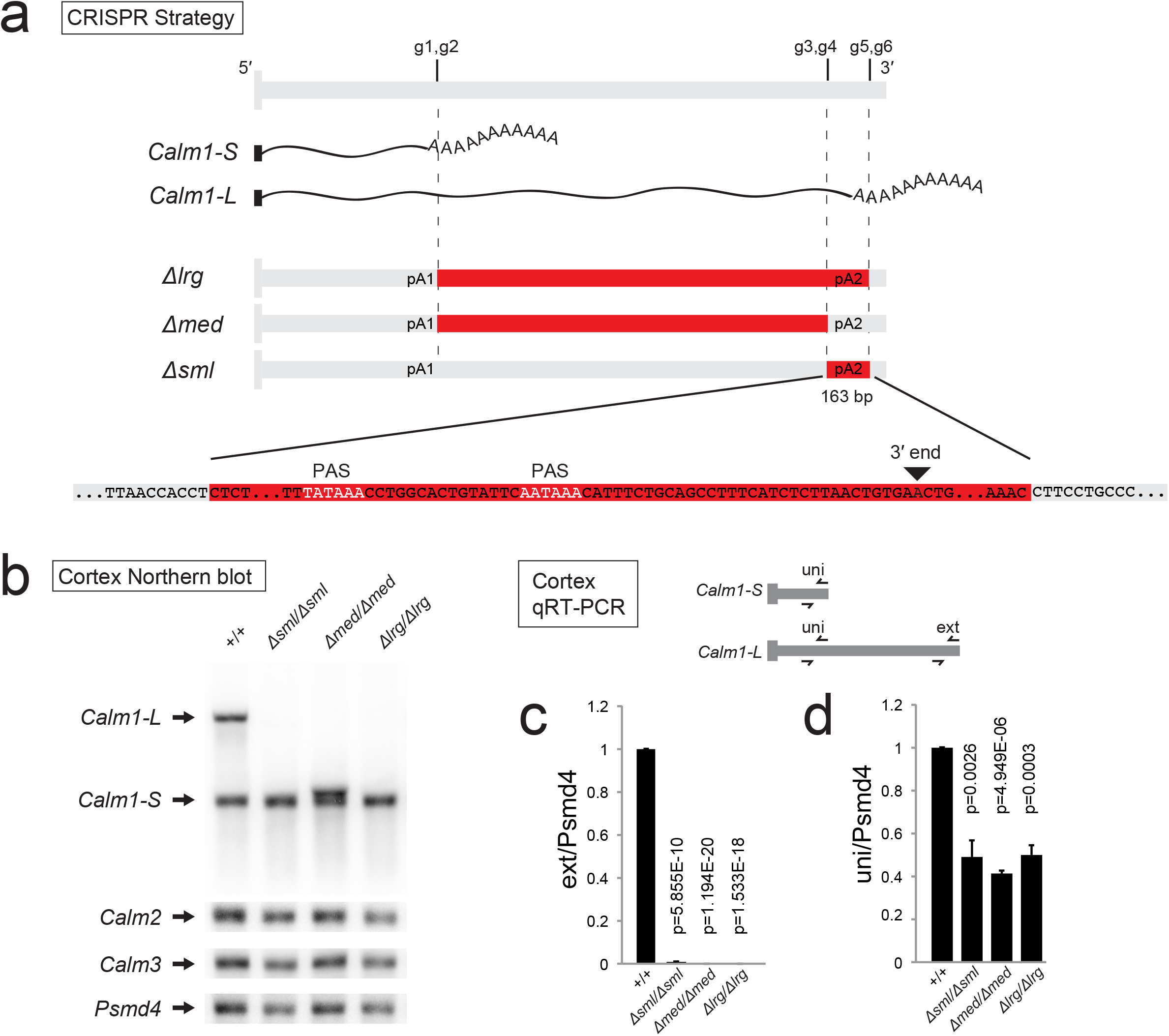
Successful generation of *Calm1* long 3′ UTR knockout mice using CRISPR-Cas9 genome editing. **(a)** Diagram of strategy used to eliminate the biogenesis of *Calm1-L*. Six gRNAs (g1-g6) were injected simultaneously to generate a variety of deletions named *Large (lrg)*, *Medium (med)*, and *Small(sml)*. Deleted regions are shown in red. Sequence removed by the *sml* deletion is shown which encompasses the distal PolyA site (pA2), including two polyA signals (PAS). **(b)** *Calm1* Northern blot of adult cortex from three deletion strains and control demonstrating successful deletion of *Calm1-L* without affecting *Calm1-S*. Note the *med* deletion generates a new isoform with a truncated long 3′ UTR due to the preservation of the distal polyA site. *Psmd4* is shown as a loading control. *Calm2* and *Calm3* levels are also shown. **(c, d)** RT-qPCR of deletion strain cortex tissues. **(c)** Long 3′ UTR specific primers (“ext”) reveal that all the deletion strains successfully prevent the production of *Calm1-L*. **(d)** Overall *Calm1* mRNA levels detected using “uni” primers was reduced approximately 2-fold when comparing wild type versus deletion cortex samples. P values from student’s t-test are shown; n=3.

Transgenic mice were generated by injection of all 6 guide RNAs along with mRNA encoding Cas9 endonuclease. From 11 founder mice we isolated 3 lines harboring different deletions that we confirmed using Sanger sequencing. One deletion, named *Large* (*lrg*), removed sequence encompassing the long 3′ UTR, including the distal polyA site. The second deletion, named *Medium (med),* deleted the majority of the sequence comprising the long 3′ UTR, save for the distal PAS. This strategy brought the proximal and distal PAS adjacent to one another, which was designed to ensure cleavage at this region and prevent selection of cryptic polyA sites further downstream. The third deletion, named *Small (sml)* was designed to remove the distal polyA site, in anticipation that although the long 3′ UTR would be transcribed, it would not be cleaved and polyadenylated, thus preventing formation of a mature *Calm1-L* transcript.

The effectiveness in preventing *Calm1-L* biogenesis in these three lines was determined by Northern blot analysis of mutant cortex samples using the universal probe. Remarkably, each deletion strategy had the desired outcome. We found that *sml* and *lrg* deletion strains had complete loss of *Calm-L* transcripts, whereas *Calm1-S* levels remained unaffected (Fig. 3b). As anticipated, Northern analysis of the *med* deletion line showed a band migrating slightly higher than the *Calm1-S* band, indicating the biogenesis of an ectopic transcript slightly longer than *Calm1-S* (Fig. 3b). Transcripts from the *Calm2* and *Calm3* genes were also monitored by Northern blot and were found to be unaltered (Fig. 3b).

To confirm the loss of *Calm1-L* using these three strategies, we performed RT-qPCR analysis. All three deletion mice were confirmed to have *Calm1-L* expression effectively eliminated using the extension primer set (Fig. 3c). The effect of the deletions on overall *Calm1* transcript levels was determined using the universal primer set, revealing that total *Calm1* levels were reduced ~2-fold in all three deletion strains compared to control (Fig. 3d). There was no difference in total *Calm1* mRNA expression when comparing among the different deletion lines.

In deciding which line to carry out phenotypic analysis, we surmised that the *med* deletion was the most confounding because an ectopic transcript was generated. The *lrg* and *sml* strategies both had the desired effect on *Calm1–L* loss without altering *Calm1-S*. We proceeded with phenotypic analysis on the *sml* allele because it had the smallest amount of genomic sequence deleted (163 bp). We reasoned that it was the least likely to alter elements such as enhancers that could be present in the long 3′ UTR-encoding region.

### Molecular characterization of Calm1^Δsml/Δsml^ mice

We hypothesized that the loss of *Calm1-L* transcript production might alter the total amount of CaM protein translated from the *Calm1* gene. However, Western analysis revealed no changes in CaM levels between wild type and mutant *Calm1^Δsml/Δsml^* adult cortex, adult hippocampus (hpc), E16.5 hindbrain, or E13.5 DRG (Fig. 4a,b). It should be noted that the CaM antibody detects protein from *Calm1, Calm2*, and *Calm3* because they produce proteins with identical amino acid sequence.

**Figure 4.**
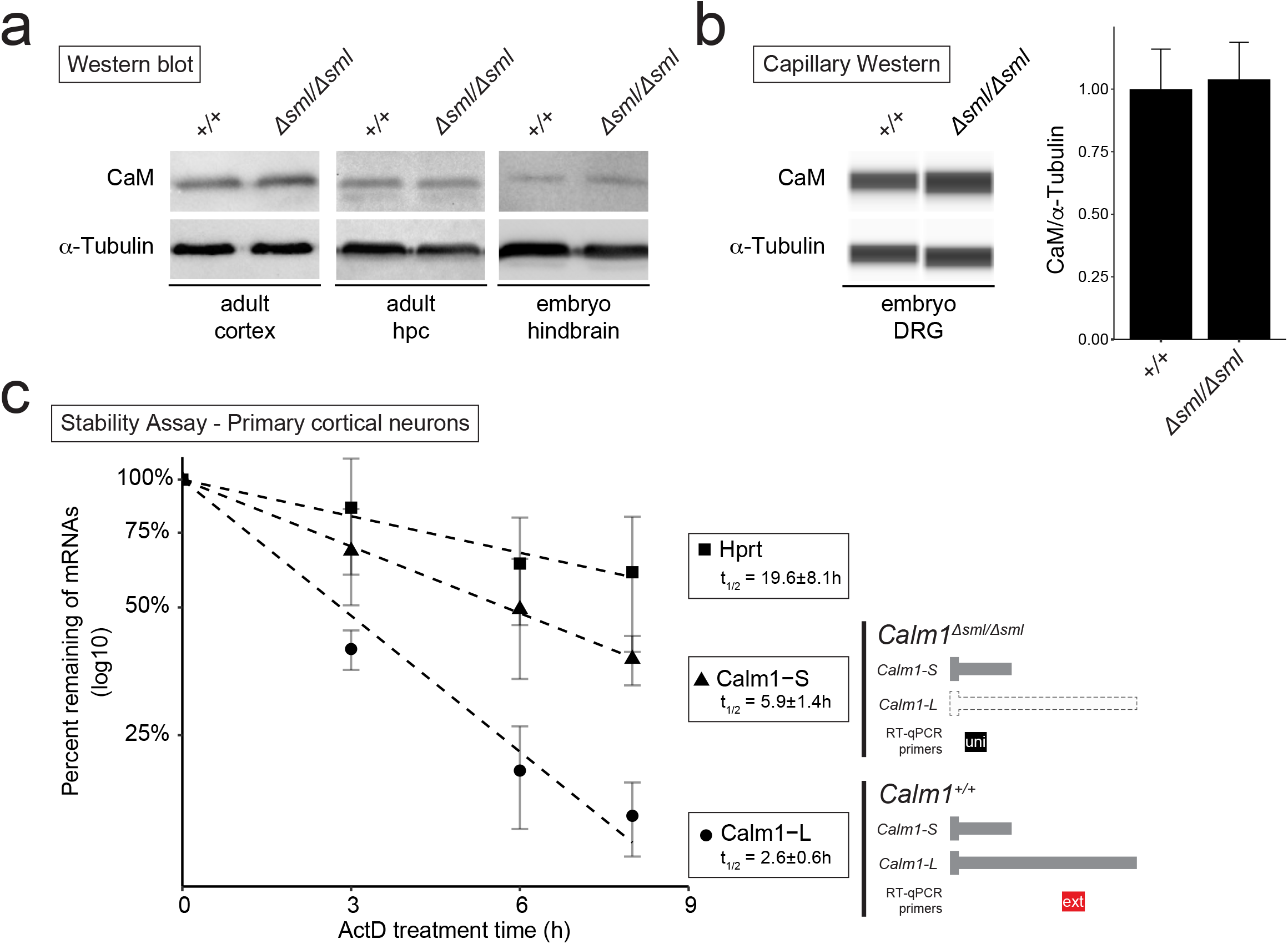
Molecular characterization of Calm1^Δsml/Δsml^ mice. **(a)** Anti-CaM western analysis of *Calm1*^+/+^ and *Calm1^Δsml/Δsml^* (*Small* deletion) samples showing no change in overall protein levels for adult cortex, adult hippocampus, or E16.5 hindbrain. **(b)** Low-input capillary western performed for E13.5 DRG. **(c)** Transcript stability assay was performed in actinomycin D treated primary cortical neurons at DIV6. *Calm1^Δsml/Δsml^* neurons were used to analyze the half-life of *Calm1-S* using the uni primer set since these neurons lack *Calm1-L* expression. The stability of *Calm1-L* isoform was assessed in *Calm1*^+/+^ neurons using the ext primer set. Hrpt mRNA was monitored as a control. This result shows that *Calm1-L* has a shorter half-life (t_1/2_= 2.6 ± 0.6 hr) compared to *Calm1-S* (t_1/2_= 5.9 ± 1.4 hr). Assay was performed using three independent biological replicates. Error bars = standard deviation of the mean.

Although we failed to detect changes in overall CaM protein levels in the mutant mice, we surmised that *Calm1-S* and *Calm1-L* should have different mRNA stabilities because of their differing 3′ UTR content. One difficulty in measuring long versus short 3’ UTR isoforms is that qRT-PCR techniques cannot uniquely detect the short isoform, due to its entire 3′ UTR sequence being shared with the long 3′ UTR. The generation of long 3′ UTR null mice presented an opportunity to employ a qRT-PCR approach for measuring the half-life of *Calm1* 3′ UTR isoforms. Primary cortical neurons from *Calm1^Δsml/Δsml^* deletion mice and *Calm1*^+/+^ controls were treated with actinomycin D for 0, 3, 6, and 8 hours, and then RNA was harvested for RT-qPCR analysis. Extension qPCR primers were used to detect *Calm1-L* from *Calm1*^+/+^ samples, and universal qPCR primers were used to uniquely detect *Calm1-S* from *Calm1^Δsml/Δsml^* samples (which do not express *Calm1-L*). An exponential regression equation was fitted to the relative abundance of each isoform from the 0 to 8 hours as relative amount = e^-k_decay_*time^ and the half-life was estimated for each transcript (Fig. 4c). A stable transcript, *Hprt*, was monitored as a control, and was found to have a half-life of 19.6 ± 8.1 hr. *Calm1-S* was found to have a longer half-life (t_1/2_= 5.9 ± 1.4 hr) compared to *Calm1-L* (t_1/2_= 2.6 ± 0.6 hr). Given that RNA stability can contribute to mRNA localization, this result is consistent with our observation of enhanced localization of *Calm1-S* versus *Calm1-L* in DRG axons (Fig. 2b,c).

### Calm1^Δsml/Δsml^ mice exhibit DRG axon development defects

*In situ* hybridization experiments had shown that *Calm1-L* expression was pronounced in the embryonic DRG (Fig. 1e). Thus, we examined *Calm1^Δsml/Δsml^* embryos for developmental defects in the DRG. E10.5 *Calm1^Δsml/Δsml^* and *Calm1*^+/+^ embryos were collected, and the DRG was imaged using anti-β3 Tubulin staining (Fig. 5a). Cell bodies of the DRG are derived from neural crest cells, which by E10.5 have already begun to form distinct ganglia adjacent to the spinal neural tube caudal to the hindbrain (Le Douarin and Smith 1988; Marmigere and Ernfors 2007). Cell bodies of the DRG send out dorsolateral axon projections concurrent with neurogenesis during this stage in development (Marmigere and Ernfors 2007). Unlike DRG that form adult structures, the first cervical (C1) DRG is a temporary embryonic population that loses its dorsolateral axonal projections and progressively undergoes programmed cell death from E10.5 to ~E12.5 (Fanarraga et al. 1997; van den Akker et al. 1999).

**Figure 5.**
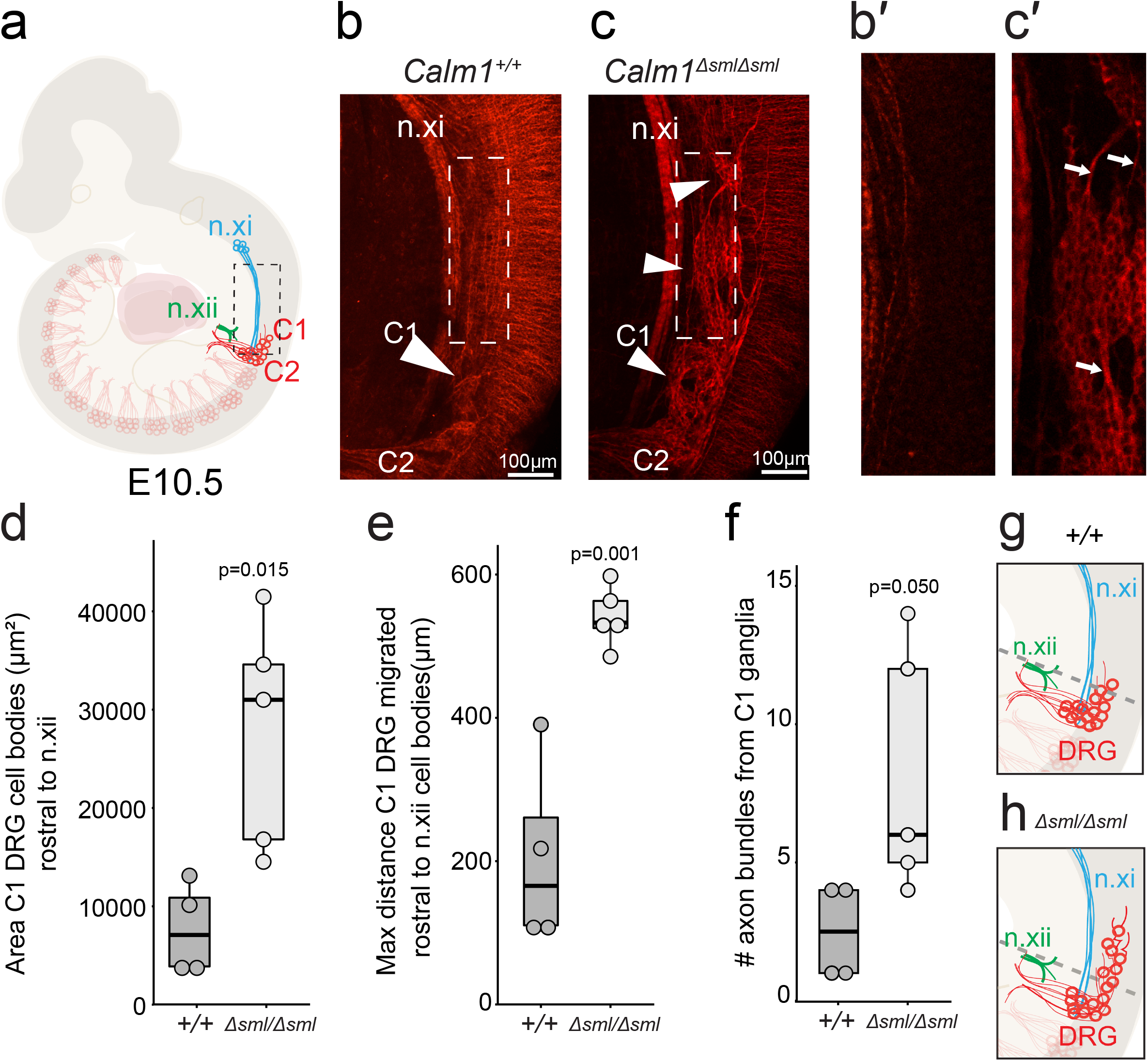
Developing C1 DRG exhibit axonal and cell body migration disorganization in *Calm1^Δsml/Δsml^* embryos. **(a)** Schematic of a wildtype E10.5 embryo highlighting the morphology of the C1 and C2 DRG axons and cell bodies. **(b, c)** *Calm1*^+/+^ and *Calm1^Δsml/Δsml^* DRG morphology visualized by anti-β3 Tubulin labeling at E10.5. **(b)** The cell bodies of the C1 DRG (arrowhead) in *Calm1^+/+^* embryos are bundled together to form a distinct ganglion. The axons of the C2 DRG can be seen projecting ventrally in an organized bundle. **(c)** The C1 DRG in *Calm1^Δsml/Δsml^* animals possesses disorganized cell bodies (arrowheads) that send out bundles of axons. Bundles of DRG cell bodies migrate rostral relative to *Calm1^+/+^* and are adjacent to the n.xi tract. **(b’, c’)** Magnified isolated section of the Z-plane highlighting regions of embryos from b) and c) focusing on disorganized mutant cell body morphology of ganglion that aberrantly migrated more rostral relative to control. **(d-h)** In order to perform measurements in the same relative location between embryos, n.xii was used as an anatomical landmark to set a beginning of measurements (indicated by the grey dashed lines in (g) and (h). **(d)** Quantification of area of ectopic cells bodies observed for developing DRG (circles represent individual data points). **(e)** Distance of aberrantly clustered cells bodies in *Calm1^Δsml/Δsml^* and *Calm1*^+/+^ (circles represent individual data points). **(f)** Quantification of the number of axon bundles projecting off C1 ganglia. **(g, h)** Representative schematic of *Calm1*^+/+^ (g) and *Calm1^Δsml/Δsml^* (h) DRG cell bodies and axons in E10.5 mouse embryos. P values determined by student’s t-test are shown; n=4 *Calm1*^+/+^; n=5 *Calm1^Δsml/Δsml^*. n.xi= accessory nerve, n.xii=hypoglossal nerve DRG= dorsal root ganglion.

The axons and cell bodies of the C1 DRG in *Calm1*^Δsml/Δsml^ embryos were found to be severely disorganized relative to *Calm1*^+/+^ embryos (Fig. 5b,c). Large groups of cell bodies of the C1 DRG in mutant embryos translocated rostral into the hindbrain adjacent to the accessory nerve (n.xi) tracks (Fig. 5c, c’) (van den Akker et al. 1999). At the same location in *Calm1*^+/+^ embryos, there were fewer translocating C1 DRG bodies that far rostral and disorganized (Fig. 5b, b’). The C1 DRG cell bodies in *Calm1*^Δsml/Δsml^ remained grouped together in smaller ganglia (Fig. 5c, c’). Axons branching off these cell bodies projected aberrantly and were tightly fasciculated (Fig. 5c’, see arrows). Some *Calm1*^Δsml/Δsml^ axons projected longitudinally, which was in contrast to the dorsolateral projections seen in non-C1 DRG (see Methods). We found significantly more C1 DRG cell bodies rostrally migrated in the mutants compared to controls (Fig. 5d). We measured the maximum distance these cell bodies migrated and found mutant cell bodies were positioned more rostral relative to controls (Fig. 5e). Lastly, we quantified the number of fascicles projecting off the C1 ganglia and found there were significantly higher numbers emanating from *Calm1*^Δsml/Δsml^ DRG compared to *Calm1*^+/+^ (Fig. 5f). Together these data suggest that *Calm1-L* is necessary for the proper development of C1 DRG axons and to restrict rostral cell migration (Fig. 5g, h).

### Explant assay demonstrates increased axon fasciculation in Calm1^Δsml/Δsml^ DRG

Previously, *Calm1* was shown to be required for NGF-dependent outgrowth of the DRG, and NGF was found to induce protein synthesis of CaM in DRG axons (Wang et al. 2015). Thus, we used recombinant NGF to test if cue-dependent outgrowth was disrupted in mutants using an *ex vivo* explant assay. E13.5 DRG tissue was plated in a collagen matrix and supplemented with or without 100ng/ml NGF for each genotype (Fig. 6).

**Figure 6.**
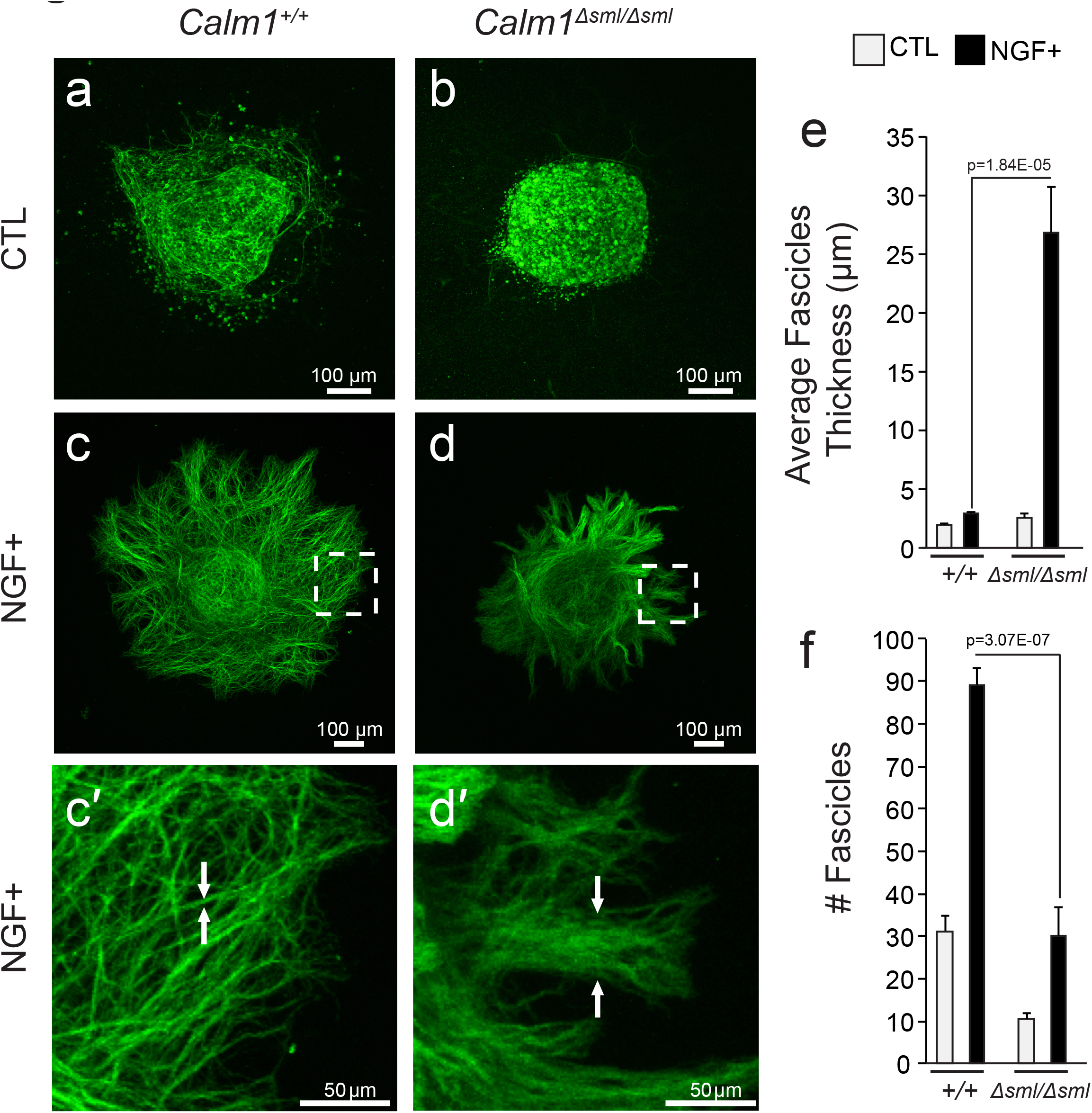
*Calm1^Δsml/Δsml^* DRG axons display increased fasciculation in explant cultures. **(a-c)** *Ex vivo* E13.5 DRG tissue visualized by anti-β3 Tubulin labeling. DRG explants grown under control conditions for both **(a)** *Calm1*^+/+^ and **(b)** *Calm1^Δsml/Δsml^* exhibit little outgrowth consisting of individual axons. **(c-d)** DRG explants treated with recombinant NGF. **(c)** In NGF treated *Calm1*^+/+^ tissue the number of axons growing off the cell bodies was increased compared to control conditions. **(d)** *Calm1^Δsml/Δsml^* DRG tissue exhibit increased numbers of fascicles growing off the explanted cell bodies when treated with NGF. *Calm1^Δsml/Δsml^* DRG explants treated with NGF possessed fascicles that were significantly larger than *Calm1*^+/+^ NGF treated DRG explants, and had overall fewer numbers of projections. **(e)** Quantification of fascicle width. NGF treated *Calm1^Δsml/Δsml^* fascicles were increased significantly compared to NGF treated *Calm1*^+/+^ fascicles. **(f)** Analysis of the number of fascicles demonstrating NGF treated *Calm1^Δsml/Δsml^* explants possess fewer numbers of fascicles overall, most likely due to their increased width. P values from student’s t-test are shown; error bars represent the standard error of the mean. n=9 untreated *Calm1*^+/+^, n=15 NGF treated *Calm1*^+/+^, n=5 untreated *Calm1^Δsml/Δsml^*, n=8 NGF treated *Calm1^Δsml/Δsml^*.

Both *Calm1*^+/+^ and *Calm1^Δsml/Δsml^* DRG neurons displayed little axon outgrowth after 48 hrs in the absence of NGF (Fig. 6a, b). In cultures with recombinant NGF added, the DRG sent out ample axonal projections for both genotypes (Fig. 6c, d). In the presence of NGF, the majority of projections in *Calm1*^+/+^ explants consisted of individual or small bundles of axons that grew out radially from the cells bodies (Fig. 6c, o’). In contrast, *Calm1^Δsml/Δsml^* tissue treated with NGF displayed axons that were fasciculated into large bundles emanating from the cell bodies (Fig. 6d, d’), which was reminiscent to bundles emanating from mutant C1 DRG *in vivo* (Fig. 5c’).

In order to quantify the degree of axon-axon interactions, we measured the overall number of fascicles radiating directly from the cell bodies in response to NGF. The average width for projections emanating from cell bodies was larger in *Calm1^Δsml/Δsml^* compared to *Calm1^+/+^* DRGs (Fig. 6e). We also counted the number of fascicles and found that there were significantly fewer fascicles projecting from the mutant DRG compared to controls (Fig. 6f). This is likely attributed to the individual bundles containing more axons. Together, these *in vivo* and *ex vivo* experiments clearly show defects in DRG development resulting from the loss of *Calm1-L*.

## Discussion

Here, we successfully implemented CRISPR-Cas9 genome editing in mice to eliminate expression of a long 3′ UTR isoform while not altering the genomic locus or expression of its corresponding short 3′ UTR isoform. To our knowledge, this is the first successful implementation of such an approach specifically for a long 3′ UTR isoform. We found that elimination of the *Calm1* long 3′ UTR isoform impaired development of the DRG in embryos, thus establishing a functional role for *Calm1-L* in neural development. Our method for long 3′ UTR deletion using CRISPR-Cas9 adds an important new tool for the characterization of these transcript isoforms which to date have few documented *in vivo* functions.

Despite the prevalence of alternative length 3′ UTR isoforms in metazoan genomes, identification of physiological functions for such transcripts using loss-of-function genetic approaches have been lacking. An *in vivo* neurological function for a long 3′ UTR isoform in mice was previously identified for the *BDNF* gene (An et al. 2008). In this study, tandem SV40 polyA sites were inserted downstream of the proximal polyA signal to prevent biogenesis of the long 3′ UTR. Our strategy for generating loss of long 3′ UTR isoforms is less confounding because artificial regulatory sequences are not inserted into the genome. Including these foreign regulatory sequences generates chimeric short 3′ UTR transcripts, which can affect cleavage and polyadenylation dynamics in unexpected ways.

We employed multiple gRNAs for CRISPR-Cas9 gene editing because it was not clear which, if any, of these strategies would effectively prevent *Calm1* long 3′ UTR biogenesis. Remarkably, a single injection of this gRNA cocktail led to three different deletions which all prevented *Calm1-L* expression. There are several advantages and disadvantages that should be considered when deciding on which deletion strategy to implement. An advantage of the *lrg* deletion strategy is that it leaves no possibility for generating the long 3′ UTR isoform. However, implementing this strategy for other long 3′ UTRs could be impractical given that many long 3′ UTR transcripts are of exceptional lengths (> 10 kb) (Miura et al. 2013) that make CRISPR-mediated deletions inefficient. Another disadvantage is that larger deletions have a greater likelihood of altering other non-coding genomic elements such as enhancers.

We were concerned that removal of the distal polyA site (as found in *sml* and *lrg* strategies) could result in the usage of downstream cryptic polyA sites that would generate unintended ectopic transcripts. Fortunately, neither the *lrg* or *sml* deletion mice showed evidence of cryptic polyA site usage. When implementing this strategy for other genes, investigators should scan downstream sequences for potential cryptic polyA signals and other *cis*-elements that promote cleavage and polyadenylation (Shi and Manley 2015).

The *med* strategy, which results in deleting the sequence in between the proximal and distal polyA sites, has been previously implemented in cell-based systems (Zhao et al. 2017). The major confounding factor for this strategy is that an ectopic, truncated mRNA slightly longer than *Calm1-S* is produced due to the distal polyA site remaining intact. Despite this drawback, this strategy might be required for some cases since bringing two adjacent polyA sites in proximity is likely to prevent usage of any downstream cryptic polyA sites.

Finally, the *sml* deletion strategy, which consisted of deleting 163 bp of sequence encompassing the polyA site, was arguably the least confounding due to the minimal deletion size. We found no evidence of cryptic polyA sites being selected in between the proximal and distal polyA sites (in the long 3′ UTR region). Nonetheless, the possibility of cryptic polyA sites becoming activated in this deletion context should be considered when implementing this strategy for other genes.

To summarize, all three deletion strategies implemented here are effective in eliminating long 3′ UTR transcripts *in vivo,* but the *sml* approach of deleting the distal polyA site is the least confounding and should be the first to consider when attempting long 3′ UTR isoform deletion. Although, we appreciate that performing CRISPR with multiple gRNAs has greater potential for more off-target effects, we see few drawbacks in choosing our approach of a single pronuclear injection with many gRNAs to generate multiple lines of mice with different 3′ UTR alleles.

Axon guidance requires execution of precise gene regulatory programs in response to extracellular cues, which results in cytoskeletal rearrangements allowing for the formation of neural circuits (Tessier-Lavigne and Goodman 1996). In recent years, gene regulation at the post-transcriptional level has been emerging as an important aspect of axon guidance (Lin and Holt 2007; Zivraj et al. 2010; Holt and Schuman 2013; Shigeoka et al. 2013). In particular, there are many examples of local translation occurring in the developing growth cone in response to extracellular cues (Campbell and Holt 2001; Lin and Holt 2007; Tcherkezian et al. 2010; Yoon et al. 2012; Preitner et al. 2016).

Our data suggests that *Calm1-L* is important for fasciculation or axon-axon contact. This was evident from *in vivo* and *ex vivo* experiments that displayed dramatically disorganized cell bodies (Fig. 5c-e), and aberrantly bundled DRG axons (Fig. 6d, d’) in *Calm1^Δsml/Δsml^* tissues. Our data also suggest that *Calm-L* is an effector of NGF-dependent regulation of axonal fasciculation. The role of NGF in DRG axon fasciculation has been long established (Rutishauser and Edelman 1980), and is dependent on Ca^2+^ levels (Tolkovsky et al. 1990). Although *Calm1-L* appears to be involved in DRG axon fasciculation, it is unclear how exactly the long 3′ UTR sequence is impacting neural development.

We have yet to determine definitively whether the loss of the long 3′ UTR leads to altered local CaM translation. We found that CaM levels were unchanged in brain tissues and DRGs of the mutant mice by Western analysis (Fig. 4a) despite the reduction of total *Calm1* transcripts (Fig. 3d). It might be that the impact of the long 3′ UTR on translational control is only seen in a spatiotemporal-specific manner. Moreover, due to the identical protein-coding sequences in *Calm1*, *Calm2*, and *Calm3* genes, we were unable to attribute any changes in CaM protein specifically to *Calm1*. Generating a mouse with an epitope tag fused to the *Calm1* coding sequence could allow for detection of CaM specifically from the *Calm1* locus. In addition, monitoring translation with methods developed to monitor protein synthesis *in situ* (tom Dieck et al. 2015) could permit visualization of altered *Calm1* translation in growing axons responding to cues such as NGF.

Upon the discovery of thousands of alternative long 3′ UTR isoforms in the brain of mouse and human, it was speculated that long 3′ UTRs might have a global role in mediating mRNA localization to dendrites and axons by gaining sequences containing mRNA localization signals (Miura et al. 2014). *Calm1* mRNA has been shown to be localized to axons and dendrites in multiple studies (Zivraj et al. 2010; Gumy et al. 2011; Kobayashi et al. 2015; Wang et al. 2015; Preitner et al. 2016; Zappulo et al. 2017; Tushev et al. 2018a). Our FISH experiments using a probe detecting all *Calm1* 3′ UTR isoforms revealed expression in cultured DRG axons. However, a probe detecting only *Calm1-L* much less signal in DRG axons, suggesting the long 3′ UTR does not promote localization in these neurons. This might not be so surprising given recent genome-wide studies of APA isoform localization patterns in neurons and neuron-like cells. Although more distal versus proximal alternative last exon APA isoforms were found to be enriched in neurites of Neuro2a cells (Taliaferro et al. 2016), a more recent study of mouse embryonic stem cell derived neurons found that short 3′ UTR isoforms were more enriched compared their long 3′ UTR counterparts in neurites compared to soma (Ciolli Mattioli et al. 2019). Another recent study found contrasting patterns for APA isoforms between soma and neuropil compartments from rat hippocampus, but no global bias for long 3′ UTR isoforms being preferentially localized to the neuropil over short 3′ UTR isoforms (Tushev et al. 2018a). High stability of the mRNA was found to correlate with neuropil localization, which is consistent with our finding here that *Calm1-S* is more stabile and axonally localized compared to *Calm1-L*.

While, the precise mechanism of how loss of *Calm1* long 3′ UTR isoform impairs DRG axon development remains unclear, our approach has nonetheless uncovered a neurodevelopment phenotype resulting from the specific loss of a long 3′ UTR isoform in mice. We have demonstrated an effective strategy for eliminating long 3′ UTR isoforms via CRISPR/Cas9 genome editing without affecting short 3′ UTR expression. This approach can now be used to determine whether other long 3′ UTR isoforms are required for neurodevelopment, and hopefully lead to new insights regarding the role of alternative long 3′ UTRs in controlling mRNA stability, localization and translation in neurons.

## Materials and Methods

### CRISPR-Cas9 genome editing of Calm1

All mice were housed in an environmentally controlled facility under the supervision of trained laboratory support personnel. Animal protocols were approved by the University of Nevada, Reno Institutional Animal Care and Use Committee (IACUC) and in accord to the standards of the National Institutes of Health Guide for the Care and Use of Laboratory Animals. The MIT CRISPR Design Tool (http://www.genome-engineering.org/crispr) was used to design guide RNAs. Guide RNAs (Supplementary Table 1–1) targeted to various regions of the *Calm1* extended 3′ UTR were cloned into the BbsI site of pX330-U6 chimeric BB-CBh-hSpCas9 plasmid (42230; Addgene). The HiScribe T7 mRNA synthesis kit (New England BioLabs) was used to *in vitro* transcribe the guide RNAs, and were subsequently purified using RNA Clean & Concentrator™-5 (Zymo Research, Cat. R1016) before assessment on an Agilent Bioanalyzer as described (Han et al. 2015).

Super-ovulating 4-6-week-old FVB/NJ female mice were mated with C57BL/6J males. After fertilization, the eggs were collected from the oviducts. Mouse zygotes were microinjected into the cytoplasm with Cas9 mRNA (100 ng/μL) and the guides RNA (100 ng/μL each) as described previously (Han et al. 2015; Oliver et al. 2015; Halim et al. 2017). After injection, zygotes were cultured in KSOM+AA medium (Millipore) for 1 h at 5% CO_2_ before transfer into the 7–10-week-old female CD1 foster mothers.

Genomic DNA was isolated from tail-snips of founder mice by overnight proteinase K digestion. Two sets of primers flanking the deletion regions were used for PCR genotyping to detect the deletion using Taq Polymerase (NEB). Sibling founder mice were crossed together to generate the F1 generation. An F1 male harboring both the *med* and *sml* alleles were crossed to C57BL/6 female mice to establish separate *sml* and *med* lines. The F1 large deletion animals were first crossed to heterozygous littermates (*lrg*/wt x *lrg*/wt). An F2 *lrg* heterozygote was crossed to C57BL/6 female mice to establish the *lrg* line. All PCR products were resolved in 1% agarose gels. To confirm the deletions and check for any insertion events Sanger sequencing was of gel purified PCR products was performed (Nevada Genomics Center, University of Nevada, Reno).

### Embryo Collection

Crossed female mice were monitored daily for vaginal plugs, with positive identification being counted as E0.5 at noon that day. Pregnant mice were euthanized at noon using CO_2_ asphyxiation, and then cervical dislocation was performed in accordance with IACUC policies at the University of Nevada, Reno. E10.5 and E13.5 embryos were extracted in PBS, with limb buds collected for genotyping, and subsequently fixed in 4% PFA in PBS. Small tears were made in the forebrain and roof plate of the hindbrain to facilitate the flow of PFA into the neural tube. Embryos were fixed by overnight incubation in 4% PFA at 4°C.

### Immunohistochemistry

For whole-mount analysis, embryos or explanted tissue were blocked and permeabilized for antibody labeling by performing 3 x 5 min, 3 x 30 min, and then overnight washes of PBST (10% FBS and 1% Triton X-100 in 1X PBS). Primary antibodies (Biolegend cat#801201 anti-β3Tubulin; 1:1000) were then added and incubated for 2 nights rocking at 4 °C. Embryos were again washed for 3 x 5 min, 3 x 30 min, and then overnight in fresh PBST. Secondary antibodies were added at a 1:200 dilution for 2 nights at 4°C rocking in the dark. Secondary antibodies were washed off again 3 x 5 min, 3 x 30 min, and then overnight in fresh PBST. Embryos were then immersed in 80% glycerol/PBS for a minimum of 2 hours, were mounted, and then imaged whole mount.

### Microscopy

A Leica SP8 TCS confocal microscope was used for imaging. Images were taken at 1048 × 1048px, 200 hz scanning speed, 3 line averaging, 8 bit, and with the use of linear-z compensation.

### DRG migration in vivo analysis

Whole mount E10.5 embryos were labeled with anti-β3Tubulin (Biolegend Cat# 801201 1:1000) and were subsequently imaged by confocal microscopy. Images analyzed using ImageJ (NIH). The cell bodies to be quantified was defined by drawing a line parallel to the last branch of the hypoglossal nerve extending across the length of the image. The hypoglossal nerve was chosen as an anatomical landmark because its morphology was unaffected in mutant embryos and served as an independent reference point so measurements were consistent across embryos. The area and distance migrated of cell bodies and axons rostral to the hypoglossal branch were measured using a freehand selection tool in ImageJ. The multipoint cell counting tool was used on ImageJ to count the number of axon bundles branching off the C1 DRG. A two-sided student’s t-test was performed to test for significance.

### Explant assay

For explant assays a collagen mixture consisting of 90 μL Rat Collagen 1 (ThermoFisher cat#A1048301), 10 μL 10X DMEM (Sigma D2429), and 3 μl of 7.5% sodium bicarbonate) was used. 20 μL of collagen mixture was pipetted in a circular pattern to create a bed of collagen at the bottom of Nunc™ 4-Well Dishes (ThermoFisher cat#144444). Collagen matrixes were allowed to solidify at room temperature for 1 hr while embryos were prepared.

E13.5 embryos were collected and dissected in Neurobasal™ Medium (ThermoFisher cat#21103049) on ice. First, a limb bud was dissected for genotyping, then abdominal organs were removed, followed by gentle disruption of the tissue adjacent to the ventral spinal cord using opened blunt forceps. A scalpel was used to remove the head at the base of the hindbrain and the caudal most end of the spinal cord to flatten out the tissue. Embryos were subsequently flipped over dorsal-side up to remove the epithelial tissue and reveal the DRG. The DRG were removed via a sharpened tungsten needle and were cleaned of any neurites projecting off and subsequently cut in half. Prepared DRG tissue was transferred to the previously prepared collagen matrix via sterilized glass pipette. Multiple pieces of tissue were placed on a single collagen bed, but were arranged to be at least ~0.1cm apart. 20 μl of collagen mixture was pipetted on top of the tissue to create a 3D collagen matrix. The collagen was allowed to solidify for an hour and was then supplemented with growth media: 50% Ham’s F12 nutrient mix (ThermoFisher cat#11765047), 40% Optimem (cat#31985062), 10% FBS, 1X Pen/Strep, 1X Glutamax (ThermoFisher cat#35050061). In NGF positive conditions, 100ng/ml of

NGF 2.5S (ThermoFisher cat#13257019) was added to the growth media. Media was replaced after 24 hrs, and cells were fixed in freshly prepared 4% PFA after a total of 48 hrs growth. Tissue was subsequently stained and imaged as described above.

### Quantification of explant axon morphology

After image acquisition, the width and number of projections was measured for explanted DRG tissue. First the images were randomly assigned numbers so analysis was performed in a blinded manner. The ImageJ line tool was used to quantify the width of fascicles emerging from the cell bodies, and the number of fascicles was determined by how many width measurements were made. Only projections immediately projecting off the cell bodies were counted. The ends of fascicles were determined by the presence of the black background flanking the projection. The mean width was calculated for each explanted tissue and used for analysis. A two-tailed Student’s t-test was used to determine significance. Standard error of the mean was used to generate error bars.

### RT-qPCR and Northern Analysis

Tissue was dissected and immediately flash frozen in liquid nitrogen and then stored at −80 °C. Tissue was subsequently pulverized using a Cellcrusher™ tissue pulverizer. RNA was then extracted using the RNeasy Plus Universal Mini Kit (Qiagen) and quantified using a NanoDrop spectrophotometer. 1μg of RNA was reverse transcribed using SuperScript III Reverse Transcriptase (Invitrogen). The 20μl cDNA reaction was diluted 5-fold in ultrapure water for use in RT-qPCR. 2μl of diluted cDNA, 0.375 μM primer concentration, 10 μl SYBR™ Select Master Mix for CFX (Applied Biosystems), in 20 μl reactions were performed in technical quadruplicate. The BioRad CFX96 real time PCR machine was used to carry out real time PCR and analysis using the delta-delta CT method was carried out using BioRad CFX Manager software. For Northern analysis, PolyA+ RNA was extracted from total RNA using NucleoTrap mRNA kit (Machery-Nagel). Northern Blot analysis was performed as previously described (Gruner et al. 2016). Briefly, polyA+ RNA samples (2 μg) was denatured in glyoxal and run in BPTE gels prior to downward transfer followed by northern blotting using 32-P dCTP labeled DNA probes (sequences of primers used to generate probes are found in Supplementary Table 1–1).

### Digoxigenin in situ hybridization

Riboprobes were generated via *in vitro* transcription using DIG RNA labeling mix (Roche) for the same probe regions as in Northern blot analysis. Sucrose cryoprotected E13.5 embryos were embedded in O.C.T compound and cryosectioned at 16 −18 μm. Sections were treated in antigen retrieval solution for 5 minutes in 95°C water bath and washed in water twice. To aid permeabilization, slides were immersed in ice-cold 20% (v/v) acetic acid for 20 sec. Upon dehydration in sequential washes in 70 – 100% ethanol, slides were stored at −20°C until use. Approximately 30 ng riboprobes were used in each hybridization at 65°C overnight. Stringent washes were performed in SSC buffer (50% formamide in 2x SSC and 0.5x SSC). Anti-DIG-AP Fab antibody (Roche) incubation was performed in 2% BSA in MABT solution for 1h at room temperature. Color development was carried out using NBT and BCIP (Roche) at room temperature until desired coloring was observed (between 16 to 20 hours). Leica DM IL LED microscope was used for imaging and stitched automatically in Leica Application Suite V4 software.

### Primary neuron culture

Cortices from postnatal day 0 (P0) – P1 mouse pups were dissected in HBSS solution, dissociated with 0.25% trypsin and plated in plating media (MEM supplemented with 0.5% w/v glucose, 0.2 mg/ml NaHCO3, 0.1 mg/ml transferrin, 10% FBS, 2 mM L-glutamine, and 0.025 mg/ml Insulin) at 73,000 cells/cm^2^ onto Poly-D-lysine coated 6-well plates. After one day in vitro (DIV), the media was replaced with growth media (MEM supplemented with 0.5% w/v glucose, 0.2 mg/ml NaHCO3, 0.1 mg/ml transferrin, 5% FBS, 0.5 mM L-glutamine, 2% v/v B-27 supplement) supplemented with 4 μM AraC (Sigma-Aldrich). The cultures were maintained until DIV6 at 37 °C with 5% CO_2_.

DRGs from E13.5 mouse embryos were dissected in Neurobasal medium. Cells were dissociated with 0.25% trypsin and triturated with a gel-loading tip. Dissociated cells were plated on PDL and laminin coated compartmentalized chamber (Xona microfluidics, XC450) or coated coverslips in culture medium (DMEM supplemented with 1X Glutamax, 10% FBS, 25 ng/ml NGF, and 1X pen/strep). In particular for the compartmentalized chamber, the somal compartment was coated only with PDL and supplemented with medium containing 10 ng/ml NGF instead of 25 ng/ml. The cultures were maintained until DIV2 at 37 °C with 5% CO_2_.

### Fluorescence in situ hybridization

Single molecular FISH was carried out in primary DRG neurons using RNAscope multiplex fluorescent v2 assay according to manufacturer’s instruction. Briefly, cells were pre-treated with permeabilization solution and protease solution; and incubated with 1X probe solution (551281-C3 for extension probe and 556541-C2 for universal probe diluted in probe diluent) for 2 hours. Subsequently, Amp1-FL to Amp4-FL were hybridized to amplify FISH signals. To combine FISH and immunofluorescence, immunostaining procedure was performed after the Amp4-FL hybridization step. Primary and secondary antibodies were incubated for 1 h at 37 °C and RT, respectively. Samples were counterstaining with DAPI and mounted in anti-fade buffer (10 mM Tris pH 8.0, 2X SSC, and 0.4% glucose in water). During image acquisition, control slide that has been incubated in probe diluent without probes were used to set laser intensity to ensure no background signal is counted in experimental conditions. Laser intensity and detector range were kept constant among different conditions. FISH-quant was used for quantification of FISH signals (Mueller et al. 2013).

### Stability assay

Cortical neurons were treated with 1 μg/ml actinomycin D at DIV6. Total RNA was collected in Trizol at 0, 3, 6, and 8 hours post-treatment. 1μg of RNA was reverse transcribed using Maxima Reverse Transcriptase (Invitrogen). qPCR was performed as described above. Expression level for each time point was calculated relative to 0 hours using BioRad CFX Manager software. Exponential regression equations were fitted for each degradation plot. Half-life of each transcript was calculated using Goal-seek function in Excel to find the time point where the relative amount of transcript is reduced to 50%. The half-life of each biological replicate was used to determine mean and standard deviation for each transcript.

### Western analysis

For conventional western blot analysis, total protein was extracted in RIPA buffer supplemented with protease inhibitor tablet (Pierce). Protein samples were separated in 15% discontinuous SDS-PAGE gel and transferred onto 0.2 μm PVDF membrane (Trans-Blot Turbo, Biorad). Membranes were blocked in 5% skim milk followed by overnight incubation with primary antibodies at 4 °C. Anti-CaM (Abcam 45689) and anti-alpha-tubulin (Sigma T9026) antibodies were used at 1:2,000 dilution. HRP conjugated secondary antibody incubation was performed at room temperature for 1 hour. HRP signal was detected using ProSignal Femto reagent (Prometheus) and imaging carried out using a ChemiDoc Touch (Biorad).

For low-input western blot analysis, capillary western system (Wes^TM^, Protein simple) was used. 500 ng of total protein lysate was loaded per capillary to visualize proteins of interest. Anti-CaM and anti-α-tubulin were used at 1:50 dilution. Quantification of band intensity was automatically estimated using Compass for SW software.

## Supplementary table legends

**Table 1:**
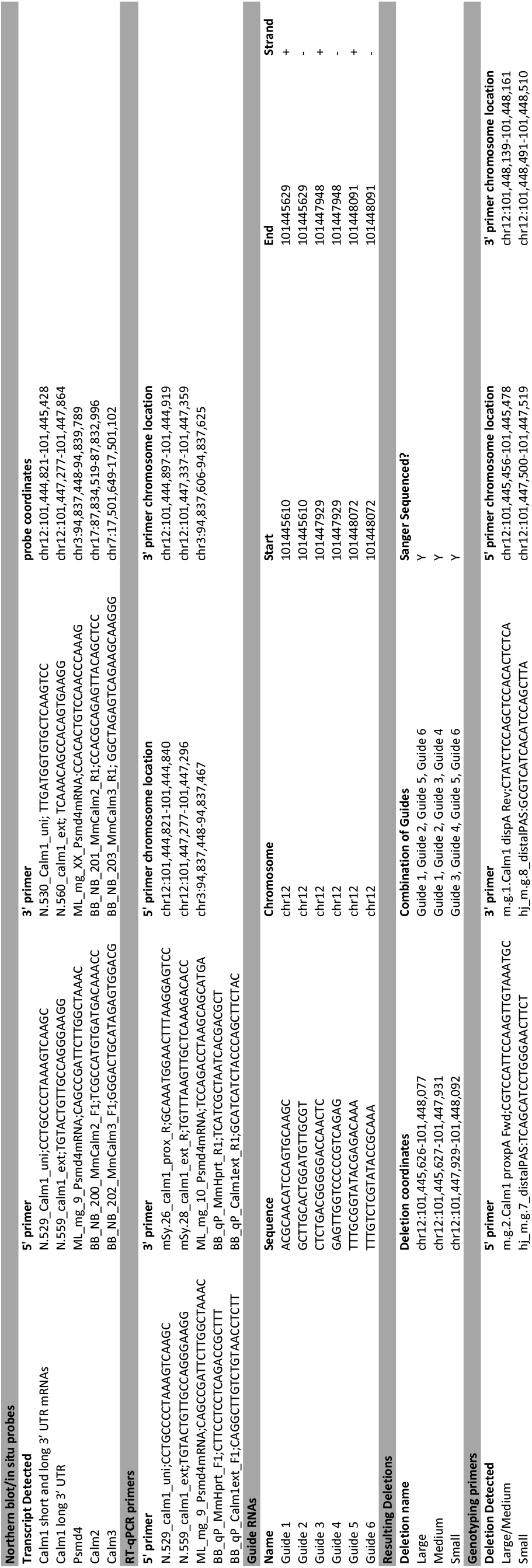
CRISPR guide sequences, deletion sequences, genotyping primers, northern blot/*in situ* probes, and RT-qPCR primers.

## Acknowledgments

We thank members of Miura lab for providing feedback and help with the project. The authors thank Vicente Gapuz III for technical assistance. We also thank Minkyung Kim and Simon Pieraut for experimental advice. Funding was provided to PM and WY from NIGMS grant P30 GM110767; GSM was supported by R01 EY025205. Core facilities at the University of Nevada, Reno campus were supported by NIGMS COBRE P30 GM103650.

